# Architecture of the cortical actomyosin network driving apical constriction in *C. elegans*

**DOI:** 10.1101/2023.01.30.526280

**Authors:** Pu Zhang, Taylor N. Medwig-Kinney, Bob Goldstein

**Affiliations:** Biology Department, University of North Carolina at Chapel Hill, Chapel Hill, North Carolina, United States of America; Lineberger Comprehensive Cancer Center, University of North Carolina at Chapel Hill, Chapel Hill, North Carolina, United States of America

## Abstract

Apical constriction is a cell shape change that drives key morphogenetic events during development, including gastrulation and neural tube formation. The forces driving apical constriction are primarily generated through the contraction of apicolateral and/or medioapical actomyosin networks. In the *Drosophila* ventral furrow, the medioapical actomyosin network has a sarcomere-like architecture, with radially polarized actin filaments and centrally enriched non-muscle myosin II and myosin activating kinase. To determine if this is a broadly conserved actin architecture driving apical constriction, we examined actomyosin architecture during *C. elegans* gastrulation, in which two endodermal precursor cells internalize from the surface of the embryo. Quantification of protein localization showed that neither the non-muscle myosin II NMY-2 nor the myosin-activating kinase MRCK-1 is enriched at the center of the apex. Further, visualization of barbed- and pointed-end capping proteins revealed that actin filaments do not exhibit radial polarization at the apex. Taken together with observations made in other organisms, our results demonstrate that diverse actomyosin architectures are used in animal cells to accomplish apical constriction.

**Summary:** Through live-cell imaging of endogenously-tagged proteins, Zhang, Medwig-Kinney, and Goldstein show that the medioapical actomyosin network driving apical constriction during *C. elegans* gastrulation is organized diffusely, in contrast to the sarcomere-like architecture previously observed in the *Drosophila* ventral furrow.

## Introduction

During development, tissues are remodeled through morphogenesis, which is largely driven by forces generated at the level of individual cells. One of the strategies utilized to achieve morphogenesis is apical constriction (Figure 1A). Apical constriction is essential for several morphogenetic events, including gastrulation in many organisms such as *Drosophila* and *C. elegans*, and neural tube formation in vertebrates (reviewed in Sawyer et al., 2010). Apical constriction is primarily driven by the contraction of actomyosin networks at the apical cortex, where non-muscle myosin II motor proteins pull on actin filaments, generating cortical tension (reviewed in Martin and Goldstein, 2014).

**Figure 1:**
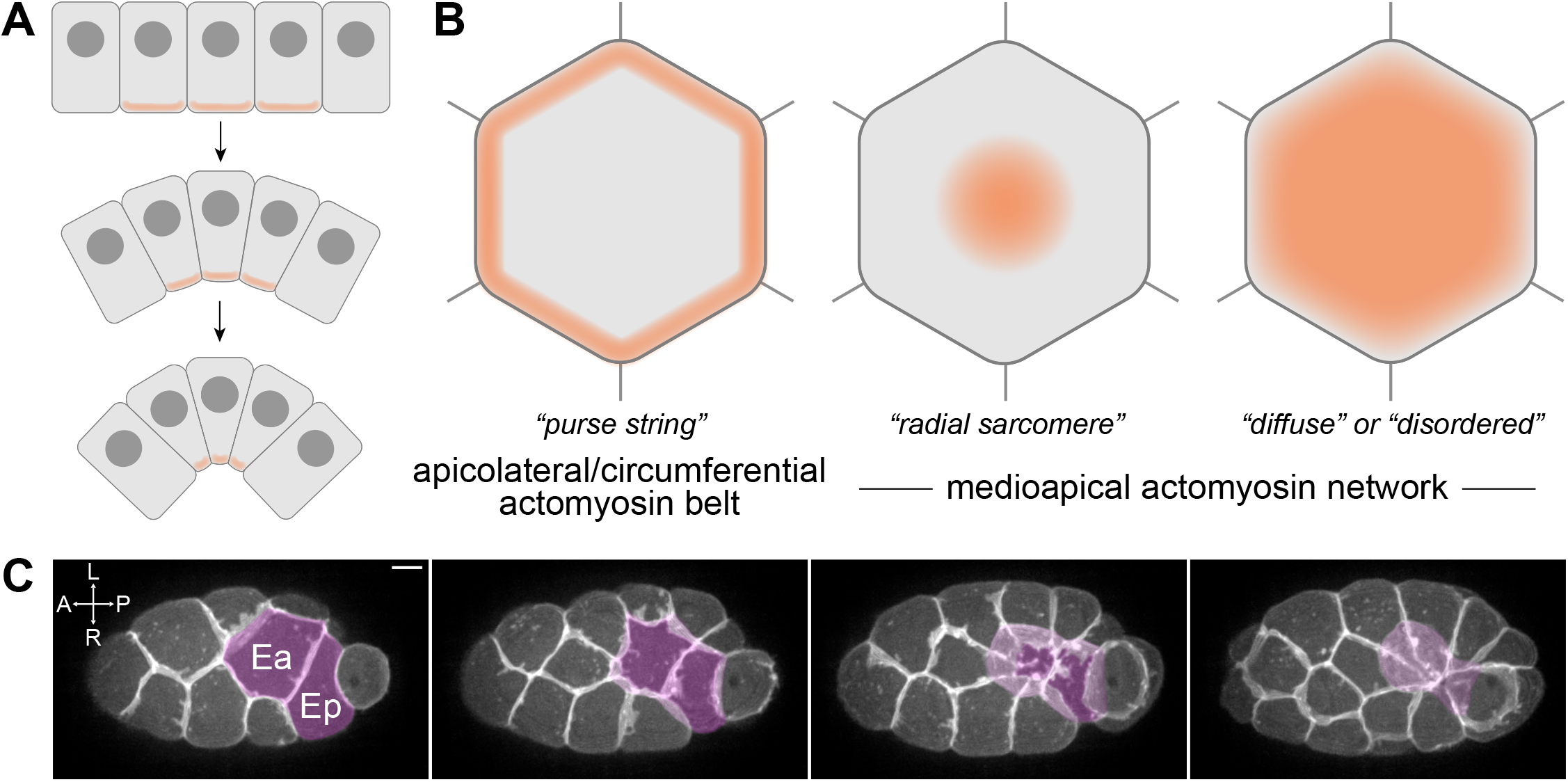
Models of apical constriction. (A) Illustration of apical constriction resulting in tissue morphogenesis. (B) Three of the models of actomyosin architecture observed in apically constricting cells. Orange shading represents the regions of localized myosin-based force generation. (C) Maximum intensity projections of 10 planes spanning 5 μm total Z-depth depicting *C. elegans* gastrulation from a ventral view with plasma membranes fluorescently labeled (*mex-5p*::mScarlet-I::PH). Ea and Ep are pseudo-colored to visualize their internalization over time. Scale bar: 5 μm.

Several models have been proposed for the organization of actomyosin networks in apically constricting cells (Figure 1B) (reviewed in Martin and Goldstein, 2014). One of the earliest described models is the circumferential contraction network or “purse string” model, which has been thought to drive apical constriction in vertebrates (Baker and Schroeder, 1967; Burnside, 1973). In this model, actin filaments and non-muscle myosin II are enriched at the periphery of the cell apex and generate pulling forces parallel to apicolateral junctions. Another model is the medioapical contractile model, in which the actomyosin network spans the cell apex and generates pulling forces largely perpendicular to apicolateral junctions (Martin and Goldstein, 2014).

The medioapical actomyosin network driving apical constriction of epithelial cells in the ventral furrow of *Drosophila* exhibits a radial sarcomere-like pattern (Coravos and Martin, 2016). In this context, actin filaments are polarized with barbed ends enriched apicolaterally and pointed ends enriched toward the center of the apex, where myosin and the myosin-activating kinase ROCK are also enriched (Coravos and Martin, 2016; Martin et al., 2009; Mason et al., 2013). Coravos and Martin (2016) found that a mutant form of ROCK that was kinase-active but distributed broadly across the medioapical cortex could disrupt apical constriction, suggesting that the normal enrichment of ROCK in the center of the apex is important in this system. This work in the *Drosophila* ventral furrow is the first to quantify the orientation of actin filaments together with the precise distributions of myosin and a myosin activator in apically constricting cells. Thus, it remains unclear to what extent other medioapical actomyosin networks that drive apical constriction share aspects of this radial sarcomere-like organization.

To determine if a radially polarized medioapical actomyosin network is a broadly conserved architecture during apical constriction, we examined actomyosin dynamics during *C. elegans* gastrulation. Gastrulation in *C. elegans* begins with the internalization of the endodermal precursor cells Ea and Ep into the embryo at the 26- to 28-cell stage (Figure 1C). This model of apical constriction is valuable for examining precise actomyosin architecture due to its amenability for live-cell imaging (Goldstein and Nance, 2020). We used live-cell imaging of endogenously-tagged proteins and quantification of protein localization to resolve the actomyosin architecture of apically constricting cells. Considering that the distribution of key proteins and orientation of actin filaments might mirror those found in *Drosophila* ventral furrow cells strongly, weakly, or not at all, we considered it important to quantitatively analyze protein localization. We report that the non-muscle myosin II NMY-2 and myosin activating kinase MRCK-1 are not enriched at the center of the cell apex, but instead exhibit punctate distribution throughout the cell apex without a central bias. Additionally, we generated reagents to visualize the polarization of actin filaments. We report that the barbed-end actin filament capping protein CAP-1 is enriched at cell-cell contacts, but the pointed-end actin filament capping protein UNC-94 is distributed throughout the cell apex, suggesting that actin filaments are not radially polarized in this system as they are in *Drosophila* ventral furrow cells.

## Results and discussion

### Non-muscle myosin II (NMY-2) is distributed throughout the apical cortex

To examine the actomyosin architecture driving apical constriction of Ea and Ep, we first assessed the localization of non-muscle myosin II using an existing strain with NMY-2 endogenously tagged with mNeonGreen (mNG) at its N-terminus (Dickinson et al., 2015). mNG::NMY-2 is enriched apically in apically-constricting cells in this strain (Marston et al., 2016; Slabodnick et al., 2022), consistent with previous observations of a fluorescently-labeled transgene (Roh-Johnson et al., 2012) and matching native protein distribution based on immunofluorescence (Nance and Priess, 2002). The mNG::NMY-2 strain is fully viable (Dickinson et al., 2015, Table S3), despite NMY-2 being an essential protein (Guo and Kemphues, 1996), indicating that N-terminally tagged NMY-2 is functional. Therefore, we used this strain to enable quantification of NMY-2 localization over time, in live embryos, to look for any enrichment in parts of the medioapical domain, and throughout the process of apical constriction. For NMY-2 and the other proteins whose distribution we quantified in this study, we collected time-lapse movies of ten or more embryos, time-aligned them using the birth of the neighboring MSxx cells as a reference point, and normalized cell lengths. We then measured intensity profiles along the anterior-posterior and left-right axes of both Ea and Ep, using Z-projections of their apical surfaces (see Materials and Methods for additional details). Given the punctate localization of mNG-NMY-2 throughout the apical cortex of Ea and Ep, we aggregated the intensity data from all samples (Figure 2C; Figure S1A). The normalized intensity plots showed no clear trend pointing to the enrichment of mNG::NMY-2 in the center of the apex of Ea and Ep (Figure 2C; Figure S1A). We conclude that non-muscle myosin II punctae are broadly distributed throughout the apical cortex of Ea and Ep, throughout the process of apical constriction.

**Figure 2:**
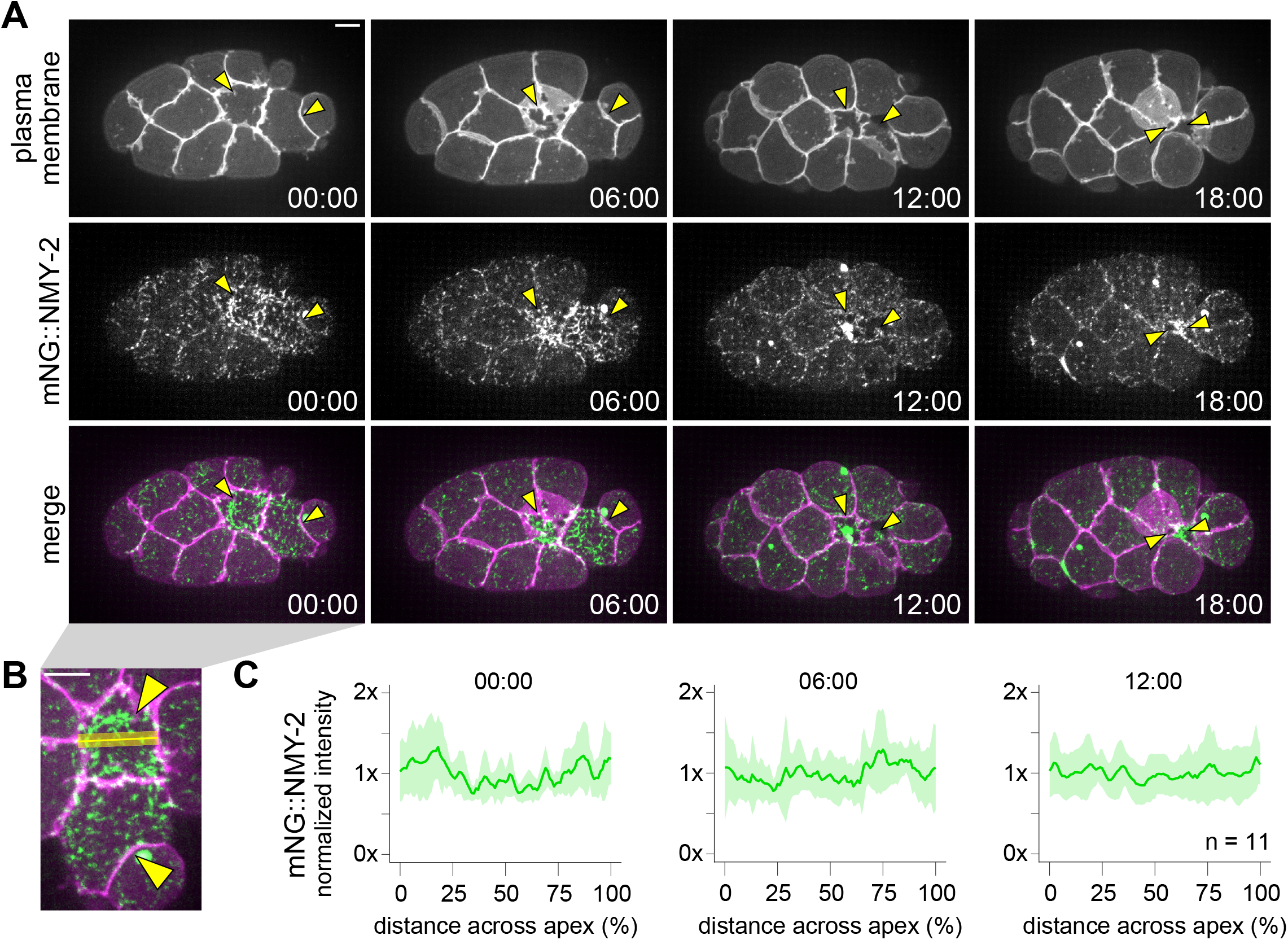
NMY-2 exhibits punctate localization distributed throughout the apical cortex during apical constriction. (A) Micrographs from a time-lapse movie depicting dynamic localization of mNeonGreen::NMY-2 over time from a ventral view. NMY-2 is co-visualized with mScarlet-I::PH, which labels the plasma membrane. Yellow arrowheads point to Ea and Ep cells. (B) An enlarged micrograph from panel A to demonstrate how line scan (yellow line) measurements were collected. (C) Plots depicting fluorescence intensity of mNeonGreen::NMY-2, normalized to the mean intensity, along the left-right axis of Ea. Solid lines indicate the mean, and shaded areas indicate the mean ± standard deviation. Time represents minutes following the birth of the neighboring MSxx cells. Scale bars: 5 μm.

### The myosin-activating kinase MRCK-1 is distributed throughout the apical cortex with slight apicolateral enrichment

While non-muscle myosin II did not exhibit enrichment at the center of the apex, it is possible that myosin activation is enriched in this region. To test this, we examined the myotonic dystrophy-related, Cdc42-binding kinase homolog, MRCK-1, which is necessary for activating myosin during Ea/Ep internalization (Marston et al., 2016). We visualized and measured localization of MRCK-1 using a YPET-tagged allele generated in preceding work (Marston et al., 2016). We found that MRCK-1 exhibits slight enrichment apicolaterally at sites of cell-cell contacts (Figure 3A,B). However, we did not find any evidence of enrichment in the center of the apex of Ea/Ep. We conclude that the myosin-activating kinase MRCK-1 is broadly distributed throughout the apical cell cortex and not enriched near the center of the apex.

**Figure 3:**
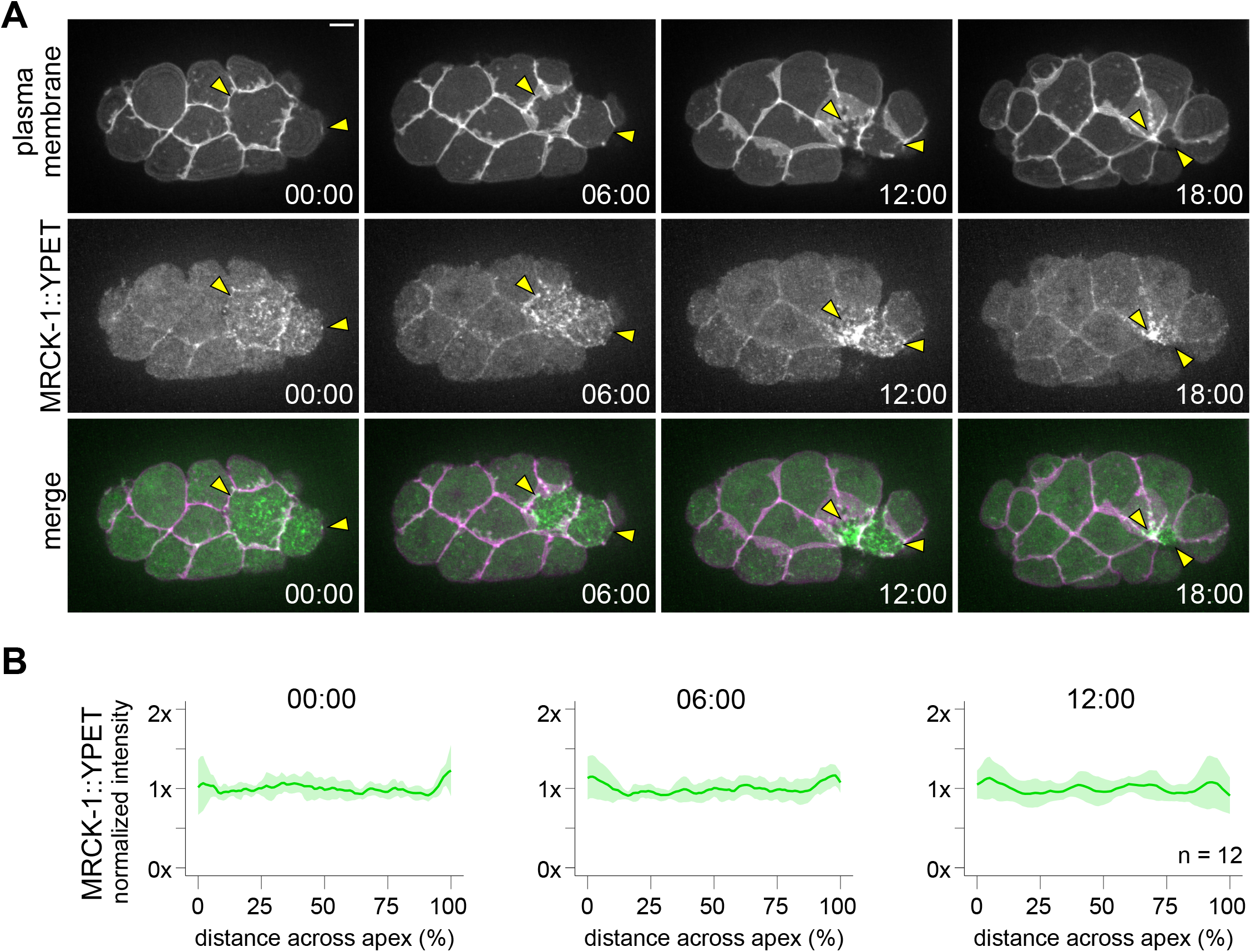
MRCK-1 exhibits slight enrichment at cell-cell borders and not at the center of the apex. (A) Micrographs from a time-lapse movie depicting dynamic localization of MRCK-1::YPET over time from a ventral view. MRCK-1 is co-visualized with mScarlet-I::PH, which labels the plasma membrane. Yellow arrowheads point to Ea and Ep. (B) Plots depicting normalized fluorescence intensity of MRCK-1::YPET along the left-right axis of Ea. Solid lines indicate the mean and shaded areas indicate the mean ± standard deviation. Time represents minutes following the birth of the neighboring MSxx cells. Scale bar: 5 μm.

### CAP-1, a barbed-end capping protein, is enriched at apicolateral junctions

Next, we sought to examine the actin architecture of the apical cortex during Ea/Ep internalization. In the *Drosophila* ventral furrow, actin filaments were observed to be organized radially with enrichment of barbed-end markers at apicolateral junctions and enrichment of a pointed-end marker in the center of the cell apex (Coravos and Martin, 2016). To visualize actin filament polarity, we endogenously tagged actin filament capping proteins. We used the gene ontology search function on WormBase (Harris et al., 2020) to identify proteins with actin filament capping functions encoded within the *C. elegans* genome, and we referenced single-cell RNA-sequencing database (Differential Expression Gene Explorer (DrEdGE)) (Tintori et al., 2016, 2020) to predict if it would be expressed at detectable levels in the embryo at the time of Ea/Ep internalization.

Using CRISPR/Cas9 genome engineering, we tagged three genes annotated as encoding proteins with barbed-end actin filament capping functions: the alpha subunit of the capping protein (CP) homolog *cap-1* (Waddle and Cooper, 1993), the receptor tyrosine kinase substrate *eps-8* (Croce et al.,2004), and the gelsolin-like protein *fli-1* (Campbell et al., 1993). The *eps-8* locus encodes twelve different isoforms due to a combination of splice variants and multiple transcriptional start sites. We tagged *eps-8* at its C-terminus with mNeonGreen as this would label all but one isoform, EPS-8B, whose essential role in the intestine has already been excluded (Croce et al., 2004) (Figure S2A). However, the sequence-confirmed EPS-8::mNeonGreen allele was expressed at levels below the detectable threshold on our imaging system, appearing similar to unlabeled, wild-type N2 control embryos at the time of Ea/Ep internalization (Figure S2B). We then tagged the C-terminus of *fli-1* and the N-terminus of *cap-1* with mNeonGreen (Figure SC-F). Both were detectable, but the *cap-1* strain was considerably brighter; thus we proceeded with *cap-1* for quantitative analyses. We retagged *cap-1* with a red fluorescent protein, mScarlet-I, in order to pair it with mNeonGreen-tagged pointed-end capping protein, described below (Figure 4A,C); the double-labeled strain was fully viable (Table S3).

**Figure 4:**
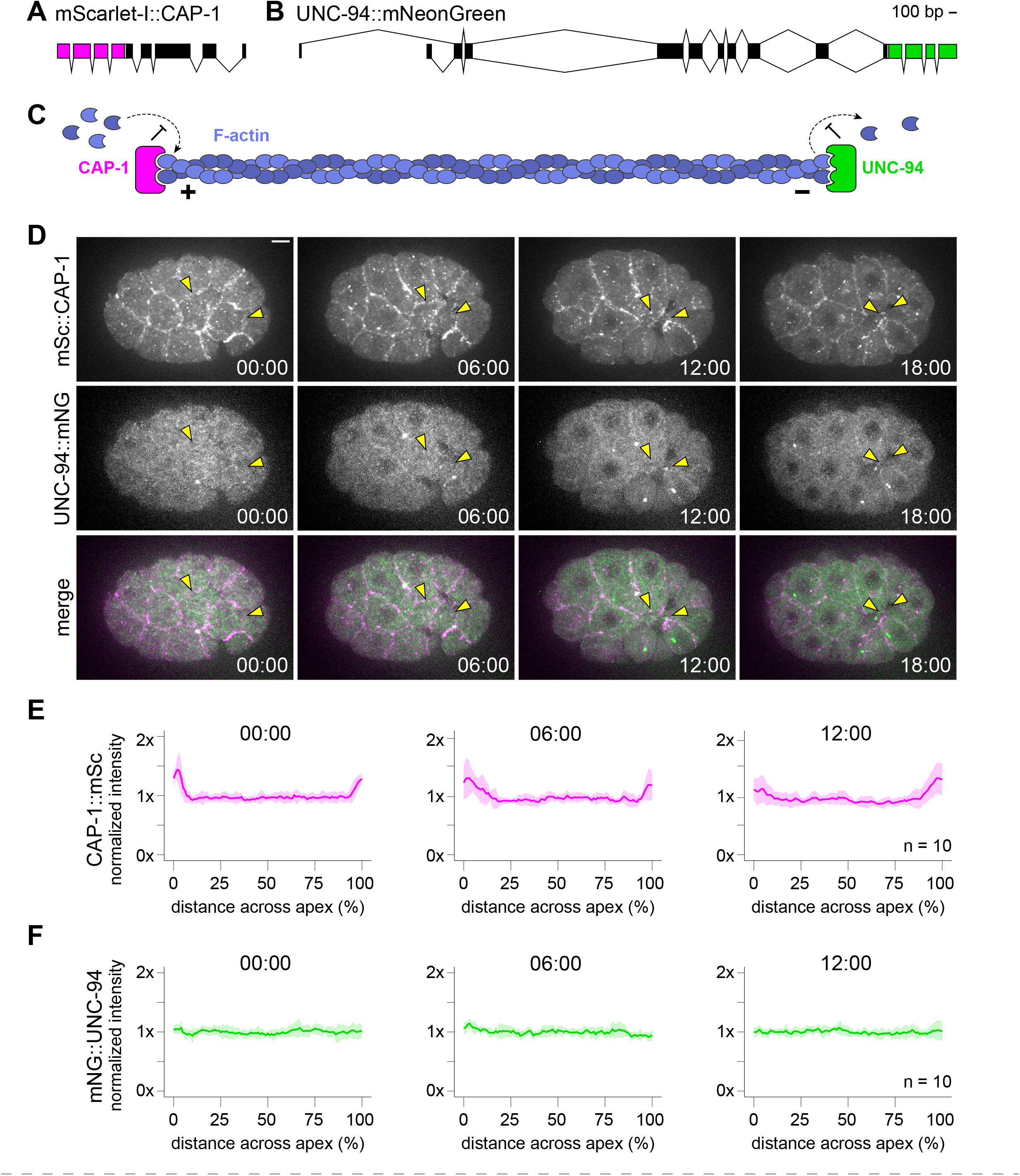
Localization of barbed- and pointed-end capping proteins, CAP-1 and UNC-94. (A) Schematic of the endogenous *cap-1* locus tagged with mScarlet-I at its N-terminus. (B) Schematic of the endogenous *unc-94* locus tagged with mNeonGreen at its C-terminus. (C) Cartoon of CAP-1 (magenta) and UNC-94 (green) performing their actin filament (blue) capping functions on the barbed (+) and pointed (-) ends, respectively. (D) Micrographs from a time-lapse movie depicting localization of mScarlet-I::CAP-1 and UNC-94::mNeonGreen over time from a ventral view. Yellow arrowheads point to Ea and Ep. (E-F) Plots depicting normalized fluorescence intensity of mScarlet-I::CAP-1 (E) and UNC-94::mNeonGreen (F) along the left-right axis of Ea. Solid lines indicate the mean, and shaded areas indicate the mean ± standard deviation. Time represents minutes following the birth of the neighboring MSxx cells. Scale bar: 5 μm.

Quantification of mScarlet-I::CAP-1 localization revealed enrichment at cell-cell contacts, with significant enrichment observed at the boundary of the Ep and P_4_ cells (Figure 4D,E, Figure S1C). This result is similar to that observed by Coravos et al. (2016) in the *Drosophila* ventral furrow, with barbed-end capping protein showing some enrichment near junctions. However, to determine if actin filaments exhibit similar radial polarity as in *Drosophila* ventral furrow cells, we also needed to examine the localization of the pointed ends.

### UNC-94, a pointed-end capping protein, exhibits is distributed throughout the apical cortex

To identify *C. elegans* genes encoding proteins with pointed-end actin filament capping functions, we again turned to the gene ontology search function on WormBase (Harris et al., 2020). Two pointed-end actin filament capping proteins were identified as homologous to tropomodulin: TMD-2 and UNC-94. According to the RNA-sequencing dataset available on DrEdGE (Tintori et al., 2016, 2020), *tmd-2* is not expressed in the internalizing Ea/Ep cells nor their precursors (E and EMS). Therefore, we focused our efforts on assessing UNC-94 localization.

The *unc-94* locus encodes two isoforms, UNC-94A and UNC-94B, with unique transcriptional start sites. This allowed us to tag each isoform independently using N-terminal mNeonGreen fusions and both isoforms together using a C-terminal mNeonGreen fusion (Figure S3A). While the mNeonGreen::UNC-94A allele was brighter than the mNeonGreen::UNC-94B allele, they exhibited similar patterns of localization (Figure S3B,C). Therefore, we utilized the C-terminally tagged version to examine the total distribution of both isoforms. Quantification of UNC-94::mNeonGreen localization revealed no evidence of enrichment in any part of the cell apex (Figure 4D,F). This result suggests that actin filaments are not organized in a sarcomere-like orientation in Ea and Ep during apical constriction.

### Concluding remarks and future perspectives

Our data suggest that the medioapical actomyosin network that drives apical constriction in Ea/Ep cells during *C. elegans* gastrulation is diffusely organized, with non-muscle myosin II (NMY-2) and myosin activating kinase (MRCK-1) distributed sparsely throughout the apical cortex. While we did observe apicolateral enrichment of the barbed-end capping protein CAP-1, its pointed-end counterpart UNC-94 did not exhibit any central enrichment. These observations are in stark contrast to actomyosin networks observed in *Drosophila* ventral furrow cells, where a radial sarcomere-like organization was found, and where there is evidence that enrichment of a myosin-activating kinase in the center of the cell apex is essential for apical constriction (Coravos and Martin, 2016). Taken together, the observations made by us and others highlight that diverse strategies of actomyosin organization are used in animal cells to accomplish apical constriction.

The radial polarization of actin filaments and centrally-enriched myosin and myosin activator seen in *Drosophila* is an organization for which contraction of the actomyosin network is a readily-expected outcome of myosin activation. Actomyosin networks with randomly oriented actin filaments might be expected to generate balanced compressive and tensile stresses, yet such networks can contract *in vitro*,owing at least in part to the buckling and severing of compressed actin filaments breaking this balance (Murrell and Gardel, 2012). Given the cortical organization that we found, we speculate that such buckling and severing might make important contributions to the forces driving apical constriction in *C. elegans*,and perhaps in other systems.

There is growing evidence that apically constricting cells rely less on the circumferential belts and more on medioapical actomyosin than initially expected (Davidson, 2012). For cells with flat or concave apical surfaces, purse-string-like contraction of circumferential belts might effectively shrink apical domains, whereas for cells with bulged apical surfaces, purse-string-like contraction of circumferential belts might be expected to pinch cells in two. In our own sampling of the literature, summarized in Table S1, we noticed apparent examples of actomyosin networks being organized medioapically in addition to circumferentially in *Xenopus* (Baldwin et al., 2022; Matsuda et al., 2022), chick (Nishimura and Takeichi,2008; Nishimura et al., 2012; Sai et al., 2014), mouse (Ebrahim et al., 2013; Lang et al., 2014; Sumigray et al., 2018; Francou et al., 2022), and the *Drosophila* salivary gland placode (Röper, 2012; Booth et al.,2014; Chung et al., 2017). It is possible that some cells and tissues use both medioapical and circumferential actomyosin networks to accomplish apical constriction. In this case, *C. elegans* gastrulation provides a valuable model to dissect the dynamics of a diffuse actomyosin network without the contribution of the circumferential networks.

With methods for CRISPR and live-cell imaging advancing, future work with other organisms may allow for precise quantification of actomyosin architecture and more cross-species comparisons. The use of endogenously-tagged proteins as done here will help avoid potential artifacts caused by overexpression in some systems, and live-cell imaging will enable the capture of protein distributions that may occur in only certain stages of apical constriction. Such information may make it possible to infer whether different actomyosin architectures are associated with other key aspects of morphogenesis, such as the internalization of few cells at a time during *C. elegans* gastrulation versus the folding of large sheets of tissue during *Drosophila* ventral furrow formation.

## Materials and methods

### Strain maintenance

The strains used in this study are listed in Table S2. Animals were maintained on NGM plates at 20 °C under standard conditions (Brenner, 1974).

### CRISPR-mediated genome engineering

The new strains presented in this work were generated through a previously described CRISPR/Cas9-mediated genome editing protocol (Ghanta and Mello, 2020). Briefly, 5 μl of 0.4 μg/μl tracrRNA and 2.8 μl of 0.4 μg/μl crRNA were added to 0.5 μl of 10 μg/μl Cas9 protein (all reagents from IDT). The mixture was incubated at 37°C for 15 minutes. 5 μl of 100 ng/μl double-stranded DNA donor with 35 bp homology arms was melted and cooled before being added to the mixture. 1.6 μl of 500 ng/μl pRF4 (*rol-6(su1006)*) plasmid was also added as a co-injection marker. Nuclease-free water was used to bring the final volume to 20 μl. The mixture was then centrifuged at 14,000 rpm for 2 minutes. 17 μl of the mixture was transferred to a fresh tube and kept on ice during the injection. Heterozygous mutants were then selected by genotyping F1 non-rollers or examining F1 under a Zeiss Axiozoom V16 stereo microscope. Homozygotes were sequenced to confirm the edits. crRNAs and homology arms sequences can be found in Table S4. Characterization of tagged alleles by comparing to known expression and localization patterns available on WormBase (Harris et al., 2020) and by measuring embryo hatching rates (Table S3). Briefly, L4-stage animals from each strain were left on 6 to 11 plates for 24 hours. The worms were then removed, and the embryos laid on each plate were counted. The embryos were examined again after 24 hours to determine the percentage that had hatched.

### Live-cell imaging

Embryos were dissected from gravid adults and mounted on slides in *C. elegans* egg buffer and sealed with VALAP. For ventral mounts, glass beads approximately 23 μm in diameter (Whitehouse Scientific Monodisperse Standards, MS0023) and clay feet were used to separate the slide and coverslip. For lateral mounts, 2.5% agar pads were used. The *mrck-1::YPET* strain was imaged with a Hamamatsu C11440-22CU ORCA Flash 4.0 sCMOS camera mounted on a Nikon Eclipse Ti inverted confocal microscope with Yokogawa QLC100/CSU10 spinning disk scan head. All other strains were imaged with a Hamamatsu ORCA QUEST qCMOS camera mounted on a Nikon Eclipse Ti inverted confocal microscope with a Yokogawa CSU-X1 spinning disk scan head. All micrographs were collected using a 60x 1.4 NA oil immersion lens and a 1.5x magnifier paired with 514 nm and 561 nm coherent lasers. Z-stacks through the ventral surface of the embryo with a step size of 0.5-1 μm were collected every 2-3 minutes.

### Image quantification

ImageJ/FIJI was used for data quantification (Schindelin et al., 2012). Intensity profiles along left-right and anterior-posterior axes were collected from maximum intensity Z-projections of the ventral surface using a line width of approximately 1.5 μm. The distance along the axis was normalized to adjust for variability in cell apex size/shape among embryos and subjected to linear interpolation sampled at each integer. The intensity was normalized to the mean. Movies were time-aligned using the timing of MSxx cell birth. The relative intensity of the mNeonGreen-tagged *unc-94* alleles was quantified from maximum intensity Z-projections of embryos at the time of MSxx cell birth. Intensity was measured by drawing a region of interest around the embryo and measuring the mean gray value, then manually subtracting the mean gray value of a background region to account for camera noise.

**Figure 5:**
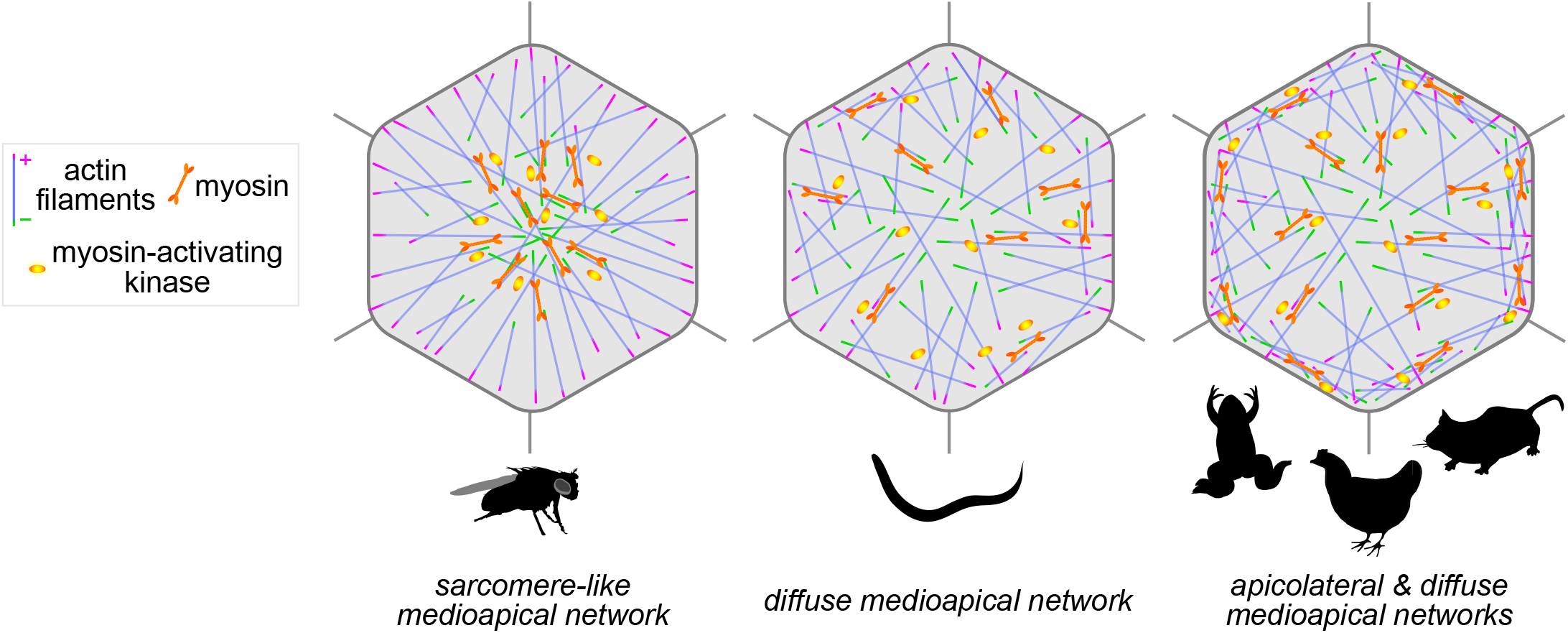
Actomyosin architectures driving apical constriction in diverse biological contexts. Illustrations depicting actin filament polarization and localization of myosin and myosin-activating kinase in the *Drosophila* ventral furrow, *C. elegans* gastrulation, and other contexts of apical constriction in *Xenopus*, chick, and mouse (see Table S1 for additional details).

## Data visualization

Representative micrographs from time-lapse movies were processed in ImageJ/FIJI (Schindelin et al.,2012). Intensity profiles were plotted using the ggplot2 package in R (Wickham, 2016). Schematics of gene loci were generated using the Exon-Intron Graphic Maker (http://wormweb.org/exonintron). Animal silhouettes were retrieved from Adobe Stock and PhyloPic (http://phylopic.org/). Figures were assembled in Adobe Illustrator.

## Acknowledgments

This work was supported by a Maximizing Investigators’ Research Award (R35GM134838 to B.G.) from the National Institutes of Health.

## Figure legends

**Figure S1:**
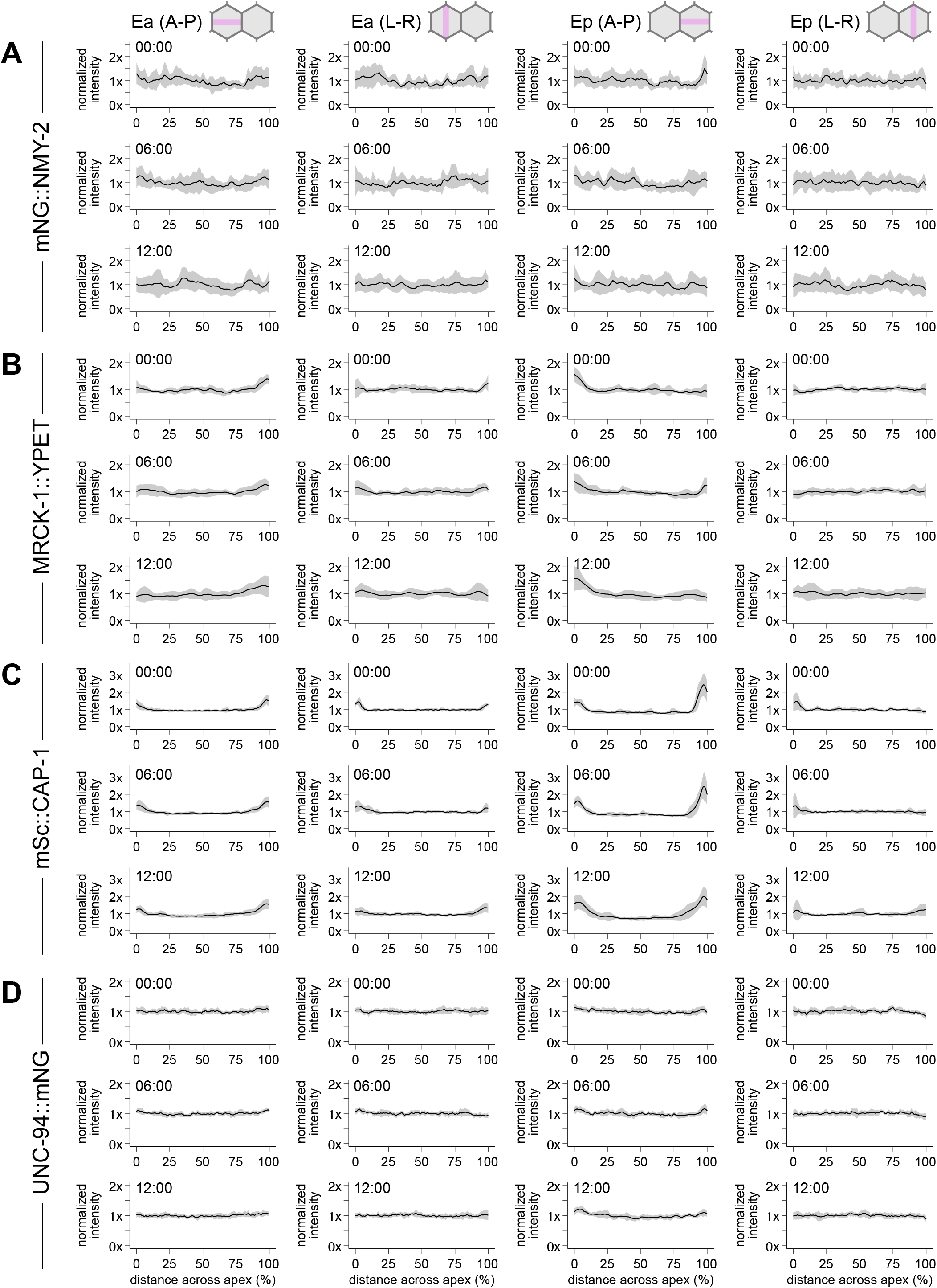
Quantification of protein localization during apical constriction. (A-D) Plots depicting normalized intensity of mNeonGreen::NMY-2 (A), MRCK-1::YPET (B), CAP-1::mScarlet-I (C), and mNeonGreen::UNC-94 (D) across the anterior-posterior (A-P) or left-right (L-R) axes of Ea and Ep. Time represents minutes following the birth of the MSxx cells. Solid lines indicate the mean, and shaded areas indicate the mean ± standard deviation.

**Figure S2:**
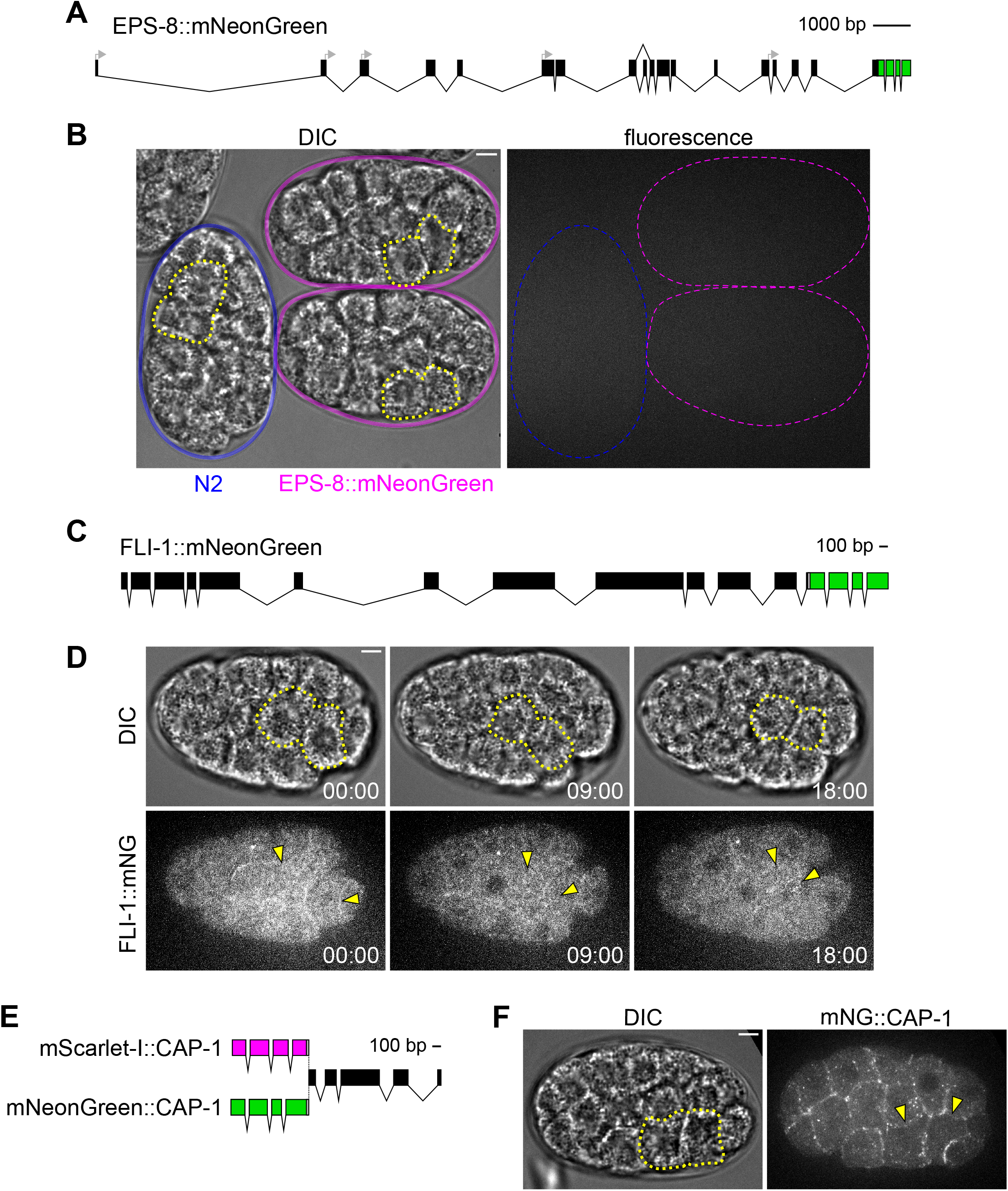
Barbed-end actin filament proteins. (A) Schematic of the endogenous *eps-8* locus tagged with mNeonGreen at its C-terminus. The start codons of the various isoforms are indicated with gray arrows. (B) DIC (left) and fluorescence (right) micrographs of N2 control (blue) and EPS-8::mNeonGreen embryos (magenta) from a lateral view. (C) Schematic of the endogenous *fli-1* locus tagged with mNeonGreen at its C-terminus. (D) DIC (top) and fluorescence (bottom) micrographs from a time-lapse movie depicting localization of FLI-1::mNeonGreen over time from a ventral view. (E) Schematic of the endogenous *cap-1* locus tagged with either mScarlet-I or mNeonGreen at its N-terminus. (F) DIC (left) and fluorescence (right) micrographs depicting localization of mNeonGreen::CAP-1 from a ventral view. Ea and Ep are indicated with yellow dotted lines or arrowheads. Time represents minutes following the birth of the MSxx cells. Scale bars: 5 μm.

**Figure S3:**
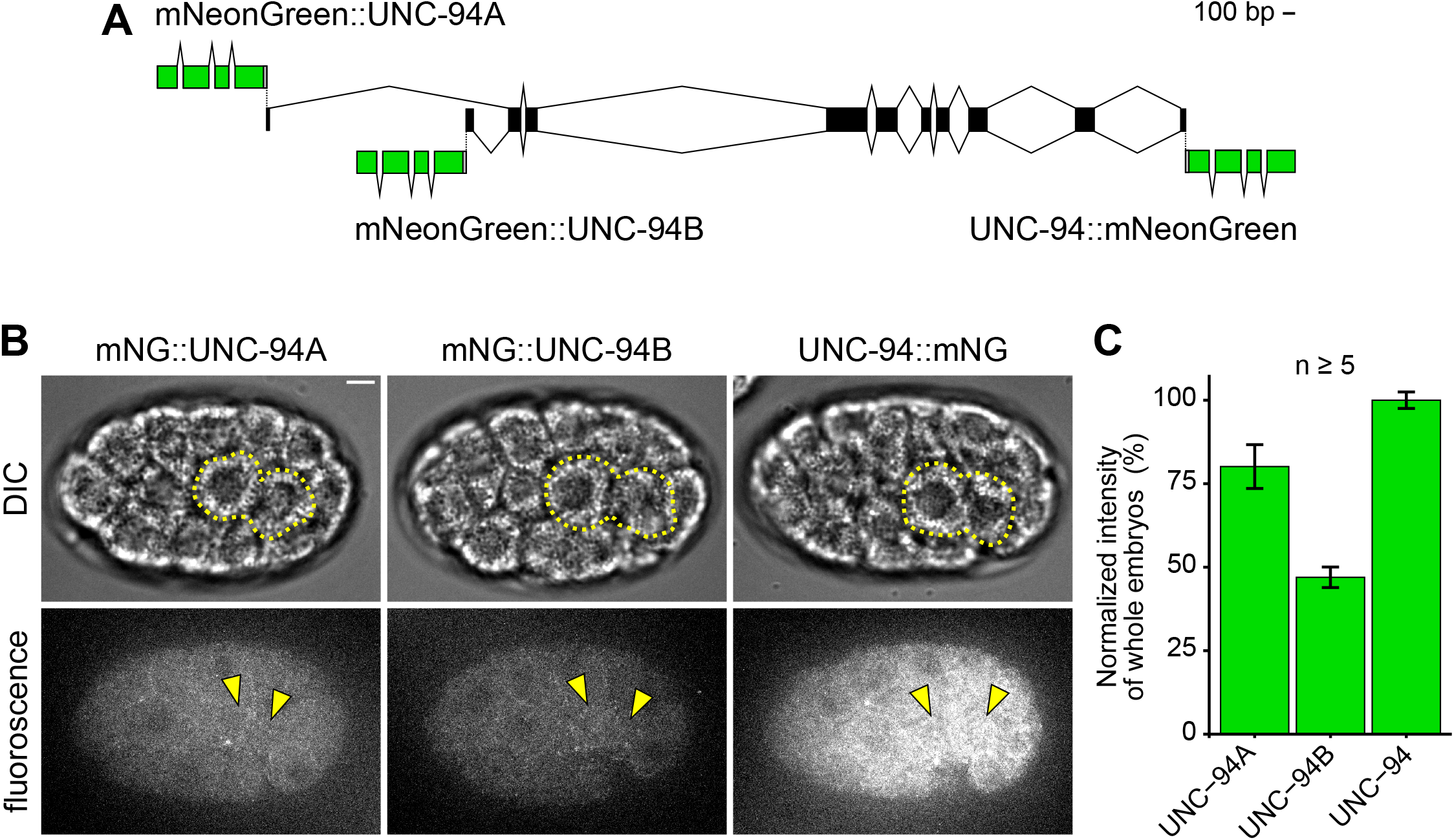
Comparison of mNeonGreen-tagged UNC-94 isoforms. (A) Schematic of the endogenous *unc-94* locus fused to mNeonGreen at its C-terminus (tagging both isoforms) or at the N-termini of the isoforms encoding either UNC-94A or UNC-94B. (B) Micrographs depicting expression and localization of UNC-94A::mNeonGreen, UNC-94B::mNeonGreen, and UNC-94::mNeonGreen from a ventral view. (C) Bar plot depicting relative intensity measurements of whole embryos with the UNC-94A::mNeonGreen, UNC-94B::mNeonGreen, and UNC-94::mNeonGreen alleles. Ea and Ep are indicated with yellow dotted lines or arrowheads. Error bars indicate the standard deviation. Scale bar: 5 μm.

## Supplementary material

**Table S1:** Literature review. Summary of our sampling of the literature reporting actomyosin organization driving apical constriction in diverse organisms and biological contexts, including our observations of published images.

| Organism | Context | Circumferential | Medio-apical | Evidence | Citation(s) |
| --- | --- | --- | --- | --- | --- |
| *C. elegans* | gastrulation |  | X | NMY-2 | Roh-Johnson et al. (2012) |
|  |  |  |  | MRCK-1 | Marston et al. (2016) |
|  |  |  |  | NMY-2 | Slabodnick et al. (2022) |
| Chick | neural tube formation | X | X | pMLC, ROCK-I | Nishimura & Takeichi (2008) |
|  |  |  |  | Myosin, pMLC, ppMLC | Nishimura et al. (2012) |
|  | otic placode | X | X | pMLC, ROCK, RhoA | Sai et al. (2014) |
| *Drosophila* | ventral furrow |  | X | Myosin | Martin et al. (2009) |
|  |  |  |  | Rok, Myosin | Mason et al. (2013) |
| | | | | Myosin, ROCK<br>Cap-α, Tmod | Coravos & Martin (2016) |
|  | salivary gland placode | X | X | Rok, Myosin | Röper (2012) |
|  |  |  |  | Myosin | Booth et al. (2014) |
|  |  |  |  | Rok, Myosin | Chung et al. (2017) |
| Mouse | organ of Corti | X | X | NMIIC | Ebrahim et al. (2013) |
|  | lens placode | X | X | Myosin IIb | Lang et al. (2014) |
|  | intestinal crypt | X | X | pMLC | Sumigray et al. (2018) |
|  | gastrulation | X | X | ppMLC, ROCK, Myosin2B | Francou et al. (2022) |
| *Xenopus* | gastrulation | X | X | myoIIb | Baldwin et al. (2022) |
|  | neural tube formation | X | X | NMIIA, pMRLC | Matsuda et al. (2022) |

**Table S2:** Strains. Names, genotypes, and associated references for the strains used in this study.

| Strain | Genotype | Source |
| --- | --- | --- |
| LP819 | *cpls131[mex-5p::mScarlet-I-C1::PH::tbb-2 3'UTR + loxN ttTi5605] II*;<br>*mrck-1(cp65[mrck-1::YPET + LoxP]) V*. | Marston et al. (2016) |
| LP823 | *nmy-2(cp132[mNG-C1^3xFlag::nmy-2]) I*;<br>*cpls131[mex-5p>mScarlet-I-C1::PH::tbb-2 3'UTR loxN ttTi5605] II*. | Slabodnick et al. (2022) |
| LP892 | *cap-1(cp436[mScarlet-I-C1::cap-1]) IV*. | This study |
| LP893 | *unc-94(cp437[mNG-C1::unc-94a]) I*. | This study |
| LP894 | *unc-94(cp438[mNG-C1::unc-94b]) I*. | This study |
| LP895 | *unc-94(cp439[unc-94::mNG-C1]) I*. | This study |
| LP896 | *unc-94(cp439[unc-94::mNG-C1]) I*;<br>*cap-1(cp436[mScarlet-I-C1::cap-1]) IV*. | This study |
| LP897 | *fli-1(cp440[fli-1::mNG-C1]) III*. | This study |
| LP898 | *eps-8(cp441[eps-8::mNG-C1]) IV*. | This study |

**Table S3:** Embryo hatching rates. Hatching rates for each of the strains used in this study.

| Strain | Genotype | Hatching rate |
| --- | --- | --- |
| LP819 | *cpls131[mex-5p::mScarlet-I-C1::PH::tbb-2 3'UTR + loxN ttTi5605] II*;<br>*mrck-1(cp65[mrck-1::YPET + LoxP]) V*. | 99% |
| LP823 | *nmy-2(cp132[mNG-C1^3xFlag::nmy-2]) I*;<br>*cpls131[mex-5p>mScarlet-I-C1::PH::tbb-2 3'UTR loxN ttTi5605] II*. | 100% |
| LP893 | *unc-94(cp437[mNG-C1::unc-94a]) I*. | 100% |
| LP894 | *unc-94(cp438[mNG-C1::unc-94b]) I*. | 100% |
| LP896 | *unc-94(cp439[unc-94::mNG-C1]) I*;<br>*cap-1(cp436[mScarlet-I-C1::cap-1]) IV*. | 100% |
| LP897 | *fli-1(cp440[fli-1::mNG-C1]) III*. | 100% |
| LP898 | *eps-8(cp441[eps-8::mNG-C1]) IV*. | 99% |

**Table S4:**
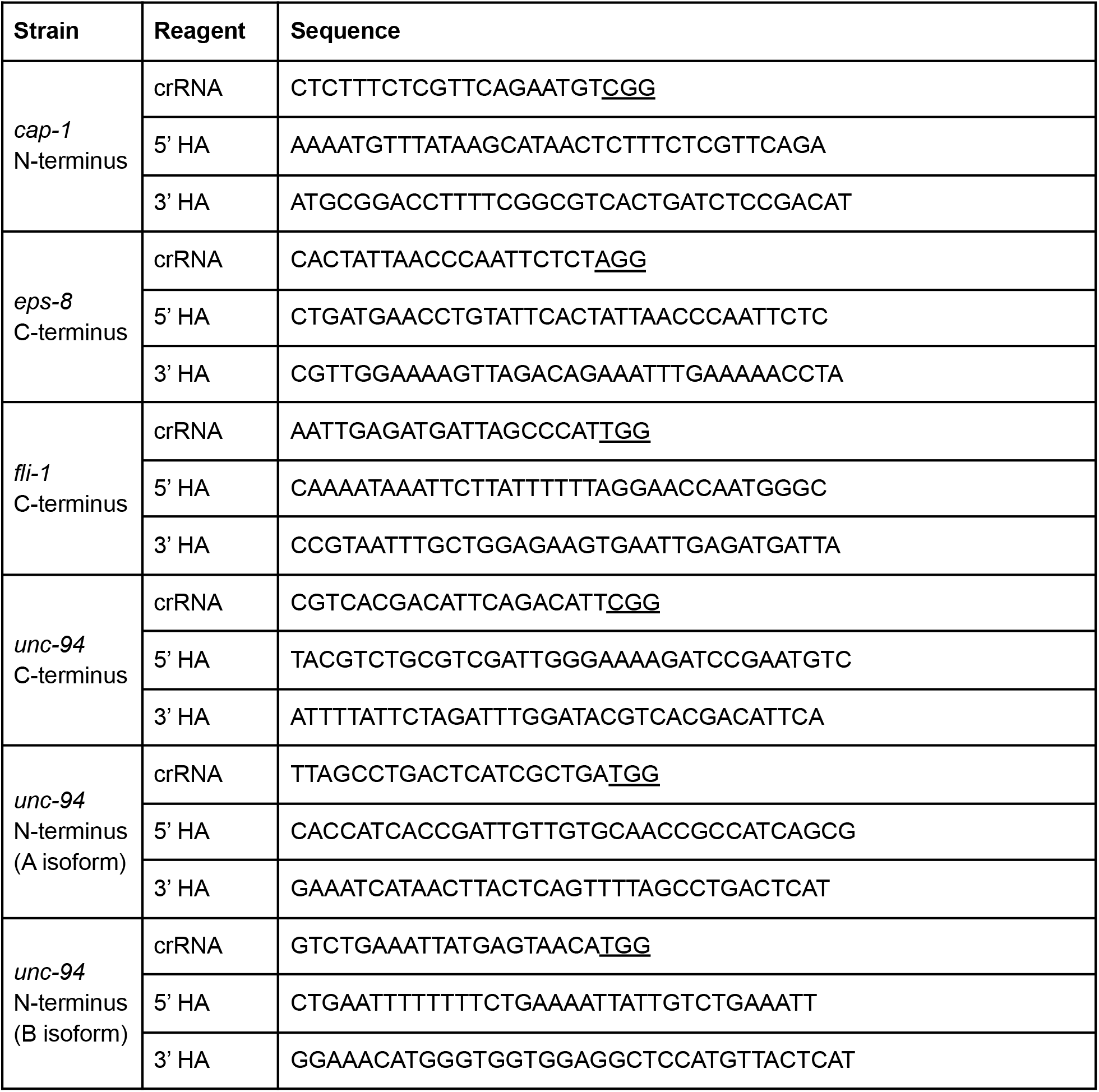
Sequences of CRISPR/Cas9 reagents. Sequences of the homology arms (HA) and CRISPR RNAs (crRNA) with PAM sequences underlined.

## Notes

### Competing Interest Statement

The authors have declared no competing interest.

